# Accurate chromosome-scale haplotype-resolved assembly of human genomes

**DOI:** 10.1101/810341

**Authors:** Shilpa Garg, Arkarachai Fungtammasan, Andrew Carroll, Mike Chou, Anthony Schmitt, Xiang Zhou, Stephen Mac, Paul Peluso, Emily Hatas, Jay Ghurye, Jared Maguire, Medhat Mahmoud, Haoyu Cheng, David Heller, Justin M. Zook, Tobias Moemke, Tobias Marschall, Fritz J. Sedlazeck, John Aach, Chen-Shan Chin, George M. Church, Heng Li

## Abstract

Haplotype-resolved or phased sequence assembly provides a complete picture of genomes and complex genetic variations. However, current phased assembly algorithms either fail to generate chromosome-scale phasing or require pedigree information, which limits their application. We present a method that leverages long accurate reads and long-range conformation data for single individuals to generate chromosome-scale phased assembly within a day. Applied to three public human genomes, PGP1, HG002 and NA12878, our method produced haplotype-resolved assemblies with contig NG50 up to 25 Mb and phased ∼99.5% of heterozygous sites to 98–99% accuracy, outperforming other approaches in terms of both contiguity and phasing completeness. We demonstrate the importance of chromosome-scale phased assemblies to discover structural variants (SVs), including thousands of new transposon insertions, and of highly polymorphic and medically important regions such as HLA and KIR. Our improved method will enable high-quality precision medicine and facilitate new studies of individual haplotype variation and population diversity.

---

Humans contain two homologous copies of every chromosome and deriving the genome sequence of each copy is essential to correctly understand allele-specific DNA methylation and gene expression, and to analyze evolution, forensics, and genetic diseases^1,2^. However, traditional de novo assembly algorithms that reconstruct genome sequences often represent the sample as a haploid genome. For a diploid genome such as the human genome, this collapsed representation results in the loss of half of heterozygous variations in the genome, may introduce assembly errors in regions diverged between haplotypes and may lead to inflated assembly for species with high heterozygosity^3^. Several algorithms have been proposed to generate haplotype-resolved assemblies (also known as phased assemblies). Early efforts such as FALCON-Unzip^4^, Supernova^5^ and our previous work^6^ use relatively short-range sequence data for phasing and can only resolve haplotypes up to several megabases for human samples. These methods are unable to phase through centromeres or long repeats. FALCON-Phase^7^, which extends FALCON-Unzip, uses Hi-C to connect phased sequence blocks and can generate longer haplotypes. Trio binning^8,9^ uses sequence reads from both parents to partition the offspring’s long reads. However, trio binning is unable to resolve regions heterozygous in all three samples in the trio and will leave such regions unphased. More importantly, parental samples are not always available, for example for samples caught in the wild or when parents are deceased. For mendelian diseases, de novo mutations in the offspring won’t be captured and phased with the parents if there are no other heterozygotes nearby. This limits the application of trio binning. Therefore, we currently lack methods that can accurately produce phased assembly for a single individual and keep pace with sequence technology innovations.

To overcome the limitations in the existing methods, we developed a new method DipAsm to accurately reconstruct the two haplotypes in a diploid individual using only PacBio’s long High-Fidelity (HiFi) reads^10^ and Hi-C data^11^ both at ∼30-fold coverage, without any pedigree information (Fig. 1). Starting with an unphased Peregrine^12^ assembly scaffolded by 3D-DNA^13^ or HiRise^14^, our pipeline calls small variants with DeepVariant^15^, phases them with WhatsHap^16^ and HapCUT2^17^, partitions the reads and assembles each partition independently with Peregrine again (Online Methods).

**Fig 1.**
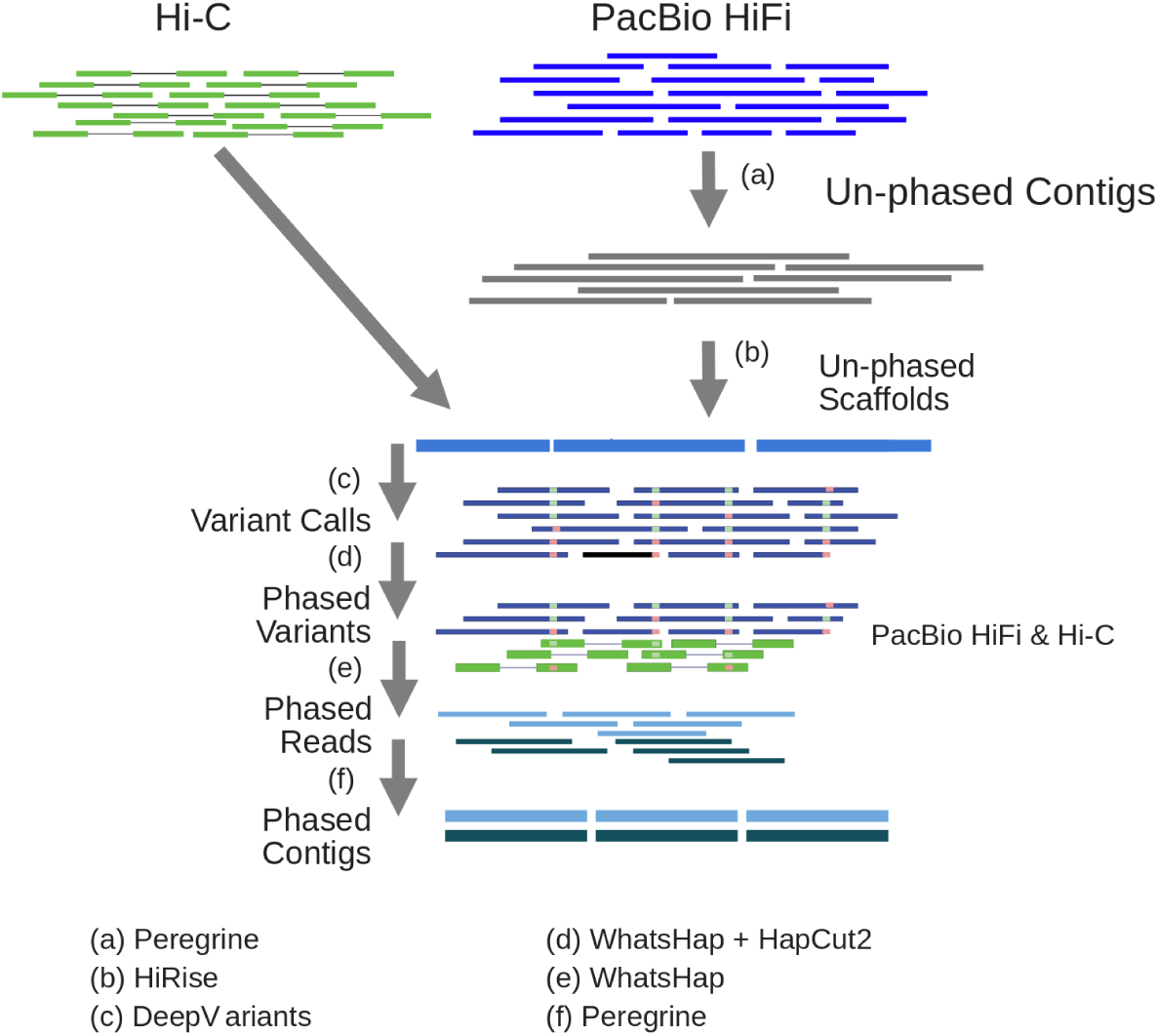
Outline of the phased assembly algorithm: DipAsm. (a) Assemble HiFi reads into unphased contigs. (b) Group and order contigs into scaffolds with Hi-C data. (c) Map HiFi reads to scaffolds and call heterozygous SNPs. (d) Phase heterozygous SNP calls with both HiFi and Hi-C data. (e) Partition reads based on their phase. (f) Assemble partitioned reads into phased contigs.

We demonstrate our method on human genomes: PGP1 from the Personal Genome Project, HG002 and NA12878 from the Genome In a Bottle dataset^18,19^ (GIAB). We produced HiFi data for the PGP1 genome and Hi-C data for HG002. (Table 1). For HG002, we also generated a trio binning based assembly with Peregrine and obtained a published Trio Canu assembly^10^ for comparison (Table 1).

**Table 1.**
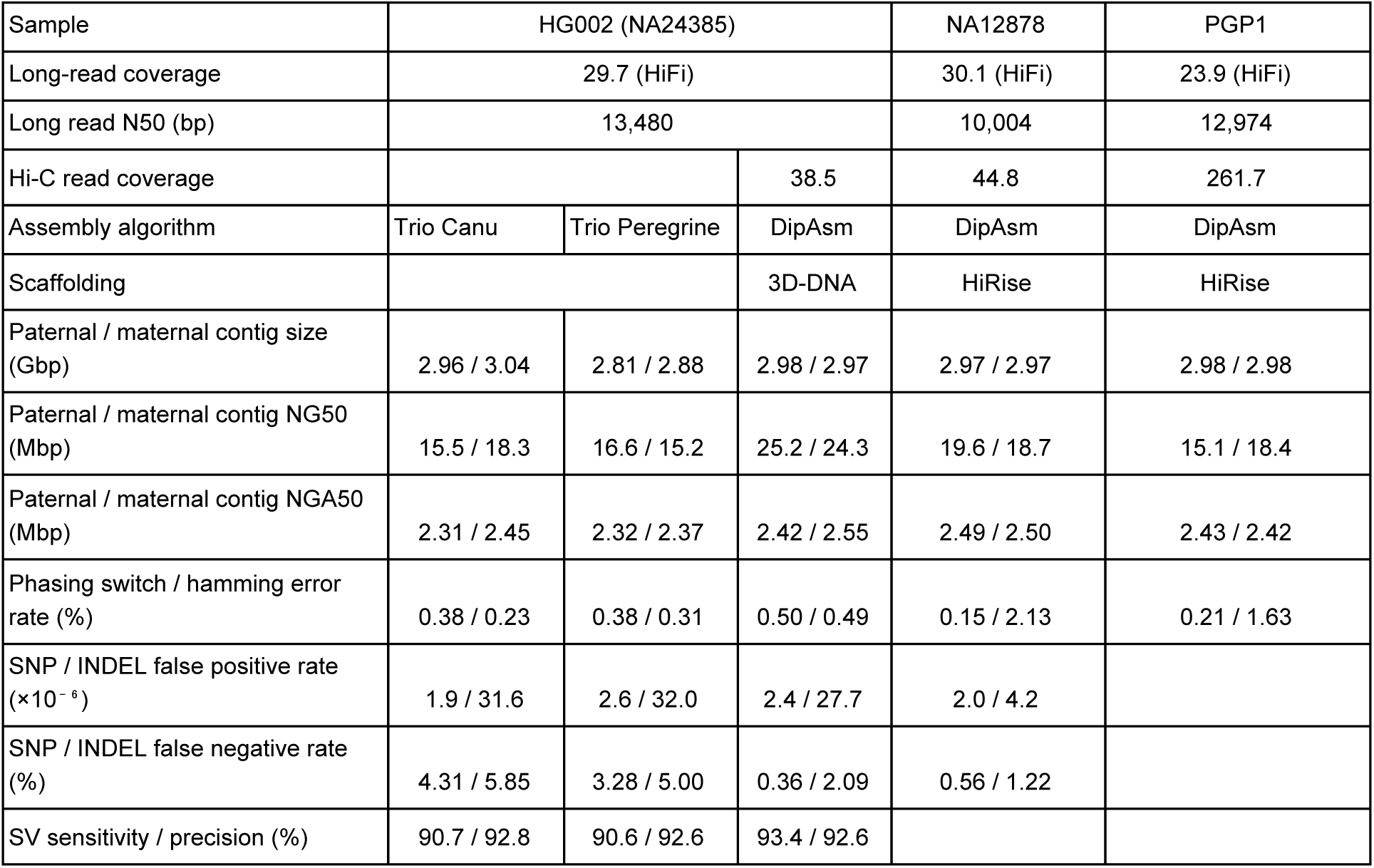
Assembly statistics. HiFi read N50: 50% of HiFi reads are longer than this number. Contig NG50: minimum contig length needed to cover 50% of the known genome (GRCh38). Contig NGA50: similar to NG50, but based on contig alignment lengths to GRCh38 instead of contig sizes. Phasing switch error rate: percent adjacent SNP pairs are wrongly phased. Phasing hamming error rate: percent SNPs wrongly phased in comparison to true phases.

From sample HG002, we generated a phased de novo assembly of 5.95 gigabases (Gb) in total, including both parental haplotypes. Half of the assembly is contained in contigs of length ∼25Mb (i.e. N50), achieving better contiguity than trio binning based assemblies. The scaffold N50 for each parent is >130 Mb. In comparison to GIAB’s SNPs phased by trio, our phasing disagrees only at 0.49% of heterozygous SNPs. This low hamming error rate over the whole genome suggests we have phased almost every chromosome into maternal and paternal haplotypes, and that the switch errors that occur only cause small local errors in phasing of a small fraction of variants.

To evaluate the consensus accuracy of our assembly, we ran the dipcall pipeline^20^ to align the phased contigs of HG002 against the human reference genome, called SNPs and short insertions and deletions (INDELs) from the alignment and then compared the assembly-based variant calls to the GIAB calls. Out of the 2.36Gb confident regions in GIAB, our de novo assembly yields 5,753 false SNP alleles (0.19% of called SNPs) and 65,302 false INDEL alleles (11.86% of called INDELs). 77% of INDEL errors are 1bp deletions, consistent with the previous observation that 1bp deletion is the major error mode for this dataset^10^. On the assumption that false positive calls are all consensus errors, not structural assembly errors or contig alignment errors, this gives a per-base error rate of 1.5×10^−5^ [=(5753+65392)/(2×2.36)] or Q48 in the Phred scale. Importantly, our de novo assembly achieves a consensus accuracy comparable to the Arrow-polished TrioCanu assembly. This suggests signal-based Arrow polishing may not be necessary for HiFi data.

The comparison to the GIAB truth data also reveals the phasing power. During assembly, failing to partition reads in heterozygous regions leads to the loss of heterozygotes and thus the elevated false negative rate in Table 1. On this metric, our Hi-C based assemblies only miss 0.4% of heterozygous SNPs, ∼8 times better than trio binning based assemblies. Trio binning is less powerful potentially because it is unable to phase a heterozygote when all individuals in a trio are heterozygous at the same site. In addition, trio binning breaks short reads into k-mers, which also reduces power in comparison to mapping full-length paired-end Hi-C reads in our pipeline. Overall this highlights even more the utility of our new approach for genetic diseases and precision medicine.

The dipcall pipeline outputs phased long INDELs along with small variants. Evaluated against the GIAB SV truth set^21^ (version 0.6) with Truvari v1.3.2, our de novo assembly based callset shows sensitivity 93.4% and precision 92.6% (Table 1). The sensitivity of trio binning based callsets is ∼3% lower, consistent with their lower sensitivity on small variants. Nearly all of the putative false positive calls are low-complexity sequences. We manually inspected some of these false positive calls from the de novo assembly. In many cases, our long INDEL calls are apparent in both HiFi read alignment and contig alignment but they are often split into multiple INDEL calls that sum to the same length as the GIAB call. Current SV benchmarking tools are unable to match SVs between vcf files when SVs are represented as multiple events in the VCF^21^. Therefore, our precision is likely substantially higher than 92.6% within the GIAB SV benchmark regions.

We additionally ran RepeatMasker^22^ on SV insertion sequences (9.1 Mb in total length) and discovered that 831, 540, and 2,303 of these are within LINEs, LTRs, and SINEs, respectively. There are 123 microsatellites, 3,582 simple repeats and 270 low-complexity sequences. We also found 21 inversions relative to the reference genome in these HG002 haplotigs (max length 25 kb, average length 5kb). A subset of SVs called from our haplotype assemblies are analyzed in Fig. 2b.

**Fig 2:**
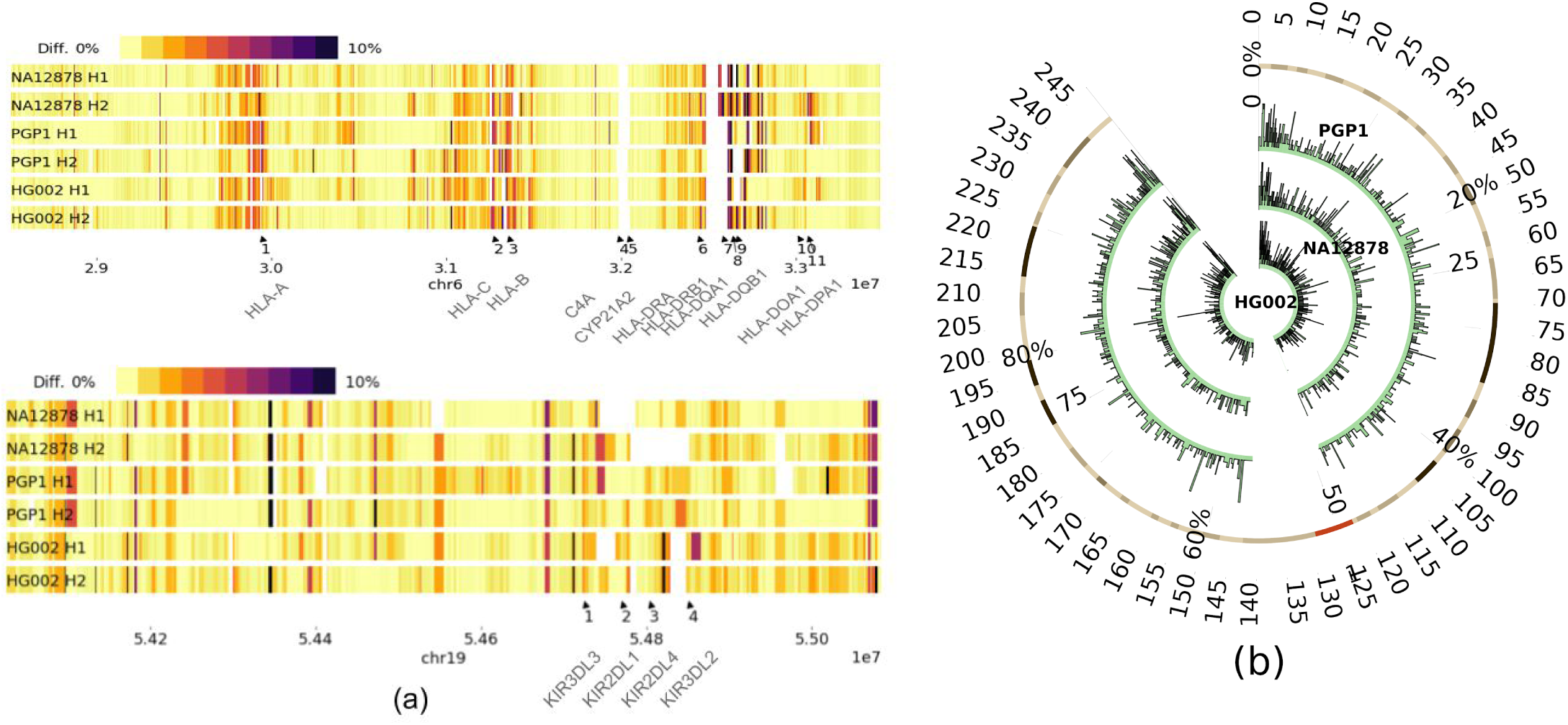
Applications of phased assemblies. (a) Local sequence divergence in comparison to the HLA (top) and the KIR (bottom) regions in GRCh38. (b) SV density per 100kb on chr1 over HG002 (inner), NA12878 (middle) and PGP1 (outer).

We assembled two other human genomes NA12878 and PGP1 with DipAsm. We can achieve chromosome-long phasing albeit the shorter read length of NA12878 and the lower read coverage of PGP1. Compared again to GIAB, the NA12878 assembly has even better consensus accuracy, measured at Q55 in GIAB’s confident regions. Interestingly, the raw HiFi base quality of NA12878 and HG002 are about the same. To understand why NA12878 has better consensus, we counted distinct 31-mers in both assemblies and HiFi reads. We found for NA12878, 3.63% of 31-mers occurring ≥3 times in reads are absent from the assembly, but for HG002, the percentage rises to 6.35%. Given that the completeness of NA12878 and HG002 are about the same, the higher percentage suggests there are more recurrent sequencing errors in HG002, which could explain the lower consensus accuracy of HG002.

The Human Leukocyte Antigen region (HLA) and the Killer-cell Immunoglobulin-like Receptor region (KIR) are among the most polymorphic regions in the human genome. Our phased assemblies can reconstruct most of these regions with two contigs for each haplotype. Based on the pattern of local sequence divergence (Fig. 2a), we can see the two haplotypes in each individual are distinct from one another. Such regions can only be faithfully assembled when we phase through the entire regions.

We present a method to generate a phased assembly for a single human individual or potentially a diploid sample of other species. It accurately produces chromosome-long phasing using only two types of input data: HiFi and Hi-C. In comparison to other single-sample phased assembly algorithms, ours is the only method capable of chromosome-long phasing. In comparison to trio binning, our method is not restricted to samples having pedigree data and can phase de novo mutations. It gives more contiguous assembly and phases a larger fraction of the genome for human samples. Meanwhile, our assembly strategy is not without limitations. First, relying on accurate SNP calls from long reads and using Peregrine for assembly, our pipeline does not work with noisy long reads at present. It is possible to switch to a noisy read assembler and to add Illumina data for SNP calling, but the assembly accuracy may be reduced due to the elevated sequencing error rate. Second, starting with an unphased assembly, we may miss highly heterozygous regions involving long SVs as demonstrated in our previous works on small genomes^6,23^. A potential solution is to retain heterozygous events in the initial assembly graph and to scaffold and dissect these events later to generate a phased assembly. Nevertheless, our improved de novo method sets a milestone. Its ability to generate phased assemblies without using a reference sequence will enable the unbiased characterization of human genome diversity and construction of a comprehensive human pangenome, which are currently goals of the Human Genome Reference Project. The ability to accurately resolve highly polymorphic regions of biological importance such as MHC and KIR, will further the goals of precision medicine.

## Acknowledgements

We are grateful to S. Koren and A. Phillippy for providing the Arrow-polished TrioCanu assembly of HG002. We thank A. English for suggesting appropriate Truvari parameters, and thank C-Z. Zhang for discussions at an early stage of this work. The authors benefited from discussions with Olga Dudchenko and the opportunity to review the results of her independent Hi-C-based phasing algorithm. This study was supported by US National Institutes of Health (grant R01HG010040 and U01HG010971 to H.L., K99HG010906 to S.G., RM1HG008525 to G.M.C. and J.A. and UM1HG008898 to F.J.S.).

## Author contributions

S.G. and G.M.C. conceived the project. S.G., C-S.C., H.L., J.A., A.F., T.Ma. and T.Mo. designed the overall strategy. S.G. implemented the assembly pipeline. M.C., E.H. and P.P. performed DNA extraction and the sequencing of PGP1 HiFi data. A.S., X.Z. and S.M. produced the HG002 Hi-C data and experimented Hi-C scaffolding with 3D-DNA. J.G. and J.M. performed the HiRise scaffolding. A.C. assisted DeepVariant calling and to improve the contig consensus accuracy. H.L., S.G., A.F., H.C., F.J.S., M.M., J.M.Z. and D.H. analyzed and evaluated the assembly. S.G. and H.L. drafted the manuscript. All authors helped to revise the draft.

## Online Methods

### PacBio CCS sequencing for PGP-1

Library Preparation: Genomic DNA was converted into a SMRTbell™ library as previously described^9^ but with a few modifications to generate slightly larger inserts. Specifically, genomic DNA was sheared using the MegaruptorR from Diagenode with the 30kb shearing protocol using a long hydropore cartridge. Prior to library preparation, the size distribution of the sheared DNA was characterized on the Agilent Femto Pulse System. A sequencing library was constructed from this sheared genomic DNA using the SMRTbell™ Template Prep Kit v 1.0 (Pacific Biosciences Ref. No. 100-259-100). In order to tighten the size distribution of the SMRTbell™ library, library was size fractionated using SageELF System from Sage Science. Approximately 4µg of SMRTbell™ Library, prepared with loading solution/Marker40. After which, the sample was loaded onto a 0.75% agarose 10kb-40kb gel cassette and size fractionated using a run target size of 7000bp set for elution well 12. A total of 8µg was fractionated on two cassettes. Fractions having the desired size distribution ranges were identified on the Agilent Femto Pulse System. Fractions centered at 11kb were pooled to generate an 11kn library and fractions centered at 16 kb were pooled to create a 16kb library. Both libraries were used for sequencing.

Sequencing: Sequencing reactions were performed on the PacBio Sequel System with the Sequel Sequencing Kit 3.0 chemistry. The samples were pre-extended without exposure to illumination for 12 hours to enable the polymerase enzymes to transition into the highly processive strand-displacing state and sequencing data was collected for 24 hours to ensure maximal yield of high-quality HiFi reads. In addition, sequencing reactions were also performed on the PacBio Sequel II System using the Sequel II Sequencing Kit 1.0 chemistry. On the Sequel II system the data collection was extended to 30 hours to ensure suitable amounts of data.

### Hi-C sequencing for HG002

A Hi-C library was generated on HG002 by Arima Genomics using a modified version of the Arima-HiC kit. Briefly, the current Arima-HiC kit (P/N: A510008) utilizes 2 restriction enzymes for simultaneous chromatin digestion. In the modified protocol, 4 restriction enzymes were deployed to enable more uniform per base coverage of the genome while maintaining the highest long-range contiguity signal, thereby benefiting analyses such as variant discovery, base polishing, scaffolding, and phasing. After the modified chromatin digestion, digested ends were labelled, proximally ligated, and then proximally-ligated DNA was purified. After the modified Arima-HiC protocol, Illumina-compatible sequencing libraries were prepared by first shearing purified Arima-HiC ligation products and then size-selecting DNA fragments using SPRI beads. The size-selected fragments containing ligation junctions were enriched using Enrichment Beads provided in the Arima-HiC kit, and converted into Illumina-compatible sequencing libraries using the Swift Accel-NGS 2S Plus kit (P/N: 21024) reagents. After adapter ligation, DNA was PCR amplified and purified using SPRI beads. The purified DNA underwent standard QC (qPCR and Bioanalyzer) and sequenced on the HiSeq X following manufacturer’s protocols.

### Phased sequence assembly

We ran Peregrine v0.1.5.2 with the following command line: “peregrine asm reads.lst 24 24 24 24 24 24 24 24 24 --with-consensus --shimmer-r 3 --best_n_ovlp 8 --output asm”, where file “reads.lst” gives the list of input read files and directory “asm” holds the output assembly. We mapped Hi-C reads to contigs with BWA-MEM v0.7.17 and scaffolded the Peregrine contigs with juicer v1.5 and 3D-DNA v180922. We preprocessed data with “juicer.sh -d juicer -p chrom.sizes -y cut-sites.txt -z contigs.fa -D”, where file “cut-sites.txt” was generated using the generate_site_positions_Arima.py script which outputs merged_nodups.txt. The scaffolds were produced with “run-asm-pipeline.sh -m haploid contigs.fa merged_nodups.txt”. We then called small variants using DeepVariant v0.8.0 with the pretrained “PACBIO” model. We mapped Hi-C reads to the scaffolds and ran HapCUT2 v1.1 over heterozygous SNP sites to obtain sparse phasing at the chromosome scale. The resulting haplotypes were then combined with PacBio HiFi data using WhatsHap v0.18 with the default parameters to generate fine-scale chromosome-long phasing. We partitioned HiFi reads based on the phases of SNPs residing on these reads, and ran Peregrine again for reads on the same haplotype from the same scaffold. This gives the final phased assembly.

With semi de novo assembly, we aligned Peregrine contigs with minimap2^24^ v2.17 against the human reference genome GRCh38 and ran RaGOO v1.1 for reference-assisted scaffolding. The phasing and re-assembly steps remain the same as in the de novo pipeline.

### Evaluating variant calling accuracy

For GIAB samples HG002 and NA12878, we compared small variant calls to GIAB v3.3.2 with RTG’s vcfeval v3.8.4. We extracted allelic errors with the “hapdip.js rtgeval” script from the syndip pipeline^20^. For sample HG002, we used Truvari v1.3.2 to evaluate long INDEL accuracy against GIAB-SV v0.6. We specified option “--passonly --multimatch” to skip filtered calls in the GIAB VCF and to allow base calls to match multiple comparison calls and vice versa. Increasing evaluation distance from the default 500 to 1000 with “-r 1000” only mildly improves the precision from 92.6% to 93.3%.

## Data availability

HG002 HiFi reads and the 250bp parental short reads were acquired from the GIAB ftp site. HG002 Hi-C reads (AC:SRR11016318), and PGP1 HiFi reads (AC:SRR11016319) sequenced by us were deposited to SRA. NA12878 HiFi reads (AC:SRX5780566), Hi-C reads (AC:SRR6675327), HG002 Hi-C reads (AC:SRR11016318), and Hi-C reads (AC:SRP173234) were downloaded from SRA. Other assemblies and assembly-based variant calls used in this work are publicly available at ftp://ftp.dfci.harvard.edu/pub/hli/whdenovo/.

## Code availability

The whole pipeline is available at https://github.com/shilpagarg/DipAsm.git

